# Lethality of leptospirosis depends on sex: male hamsters succumb to infection with lower doses of pathogenic Leptospira

**DOI:** 10.1101/322784

**Authors:** Charles K. Gomes, Mariana Guedes, Hari-Hara Potula, Odir A. Dellagostin, Maria Gomes-Solecki

## Abstract

Leptospirosis is a widespread zoonotic disease caused by pathogenic spirochetes of the genus *Leptospira* which affect both humans and animals. A somewhat contradictory published body of evidence suggests that sex impacts severity outcomes of human Leptospirosis. In this study, we used an acute animal model of disease to analyze how male and female hamsters infected side-by-side with low but increasing doses of *L. interrogans* under the same exposure conditions develop Leptospirosis. We found that female hamsters were considerably more resistant to Leptospirosis given that 87.5% survived infection; male hamsters did not gain weight and 93.7% succumbed to infection with the same infectious doses. Analysis of bacterial burden in kidney of male hamsters showed that infection with the lowest dose (10^3^) resulted in a 4Log increase of *L. interrogans*, whereas females infected with the same dose had a reduction of ~1Log, after 28 days of infection. Non-surviving hamsters had signs of compromised renal function (higher levels of creatinine in blood), as well as increased levels of anti-inflammatory IL-10 and innate response pro-inflammatory CCL3, CxCL10 and TNF-α in kidney, as well as ColA1 which is a marker of kidney fibrosis. In endemic areas, humans are more often exposed to the lower infection doses which were lethal to males, than the higher doses which were lethal to males and females. It is possible that the frequency of lower infectious doses in nature and increased biological susceptibility both contribute for aggravated outcomes of leptospirosis in males.

## INTRODUCTION

Leptospirosis is an emerging widespread zoonotic disease caused by pathogenic spirochetes of the genus *Leptospira*. The infection commonly occurs through direct contact with infected urine or indirectly through contaminated water (1). Caused by over 250 serovars of *Leptospira* spp., which are mainly hosted by rodents, leptospirosis shows clinical manifestations in humans that vary from flu-like symptoms to multi-organ failure (2). In humans, morbidity estimates are ~1 million cases a year worldwide with a 5-10% mortality rate (1), (3).

Differences in susceptibility to inflammatory responses between males and females have been noted for a number of years and sex is now accepted as a risk factor to infectious and autoimmune diseases (4), (5), (6). Although evidence that women are more susceptible to leptospirosis was reported in the past (7), more recent clinical and sero-epidemiological evidence suggests that incidence of human Leptospirosis is higher in males than in female adults and children (8), (9), (10), (11). Furthermore, its severe clinical signs requiring hospitalization are also more frequently observed in men (12), (13). Most studies focused on identification of motility factors, LPS biosynthesis and outer membrane proteins using animal models of acute Leptospirosis have been done using male hamsters (14), (15), (16), (17), (18), (19), (20); few used females and one investigator reports the use of both sexes (21), (22), (23). The US National Institutes of Health (NIH) established new guidelines to enhance reproducibility of scientific results that mandate the analysis of the effect of sex differences in cell and animal studies (24), (25). The goal of our study was to infect male and female hamsters side-by-side with low but increasing doses of the same strain of *L. interrogans* ser. Copenhageni FioCruz and analyze differences in pathophysiology and disease severity.

## MATERIAL AND METHODS

### Animals and ethics statement

Adult male and female Golden Syrian hamsters (*Mesocricetus auratus*), n=40, 8 weeks old, were obtained from Charles River Laboratory. The animals were housed in groups of 2 per cage (same sex) in an ABSL-2 pathogen-free environment in the Laboratory Animal Care Unit of the University of Tennessee Health Science Center (UTHSC). This study was carried out in accordance with the Guide for the Care and Use of Laboratory Animals of the NIH. The protocol was approved by the UTHSC Institutional Animal Care and Use Committee, Animal Care Protocol Application (Permit Number: 16-070).

### Bacterial strains and culture

*Leptospira interrogans* serovar Copenhageni strain Fiocruz L1-130 was cultivated in Ellinghausen-McCullough-Johnson-Harris (EMJH) medium supplemented with BD™ Difco™ Leptospira enrichment EMJH at 30°C. The culture (passage 4) was allowed to reach log phase of growth and pelleted by centrifugation at 3000 × g for 5 minutes, washed and resuspended in sterile 1X PBS. The cells were counted under dark-field microscopy (Zeiss USA, NY) using a Petroff-Hausser chamber.

### Experimental infection

Overall, 20 male and 20 female hamsters were housed in same sex pairs. Groups of animals were injected intraperitoneally with 10^3^ (n=8), 5×10^3^ (n=8), or 10^4^ (n=16) *Leptospira* in 1 mL 1X PBS. Non-infected (Controls) animals (n=8) were injected with an equal volume of sterile 1X PBS. Survival and body weight were monitored for 28 days post-infection given that weight loss has been found to be an early objective sign of clinical leptospirosis. Animals were scored for signs of clinical illness (body weight loss > 10%, loss of interest in food or water, prostration, ruffled fur) and were euthanized by isoflurane overdose when they reached the endpoint criteria, or at 28 days post infection. Blood from euthanized animals was collected in EDTA by cardiac puncture. After dissection, one of the kidneys was collected and stored in 1 mL RNAlater (Sigma-Aldrich) for molecular bioassays and the other kidney was placed in EMJH media for culture of *Leptospira* (as in (26)).

### q-PCR and RT-PCR

DNA was extracted from kidney using a tissue kit (NucleoSpin). *Leptospira* was quantified using TAMRA probe and primers (Eurofins) to *Leptospira* 16s rRNA by qPCR. A standard curve obtained from serial 5-fold dilutions of known numbers of *Leptospira* was used for absolute quantification. Results were expressed as the number of *Leptospira* equivalent genomes per mg of kidney tissue DNA. Total RNA was extracted from kidney tissues using a RNeasy mini kit and transcribed using a high-capacity cDNA reverse transcription kit. cDNA was subjected to real-time PCR using SYBR Green and primers as previously described (27), (28). PCR data are reported as the relative increase in mRNA transcript levels of MIP1-α (CCL3), IP-10 (CXCL10), TNF-α, IFN-γ, IL-4, IL-6, IL-10, iNOS and ColA1 of male and female hamsters infected with *L. interrogans* using β-actin as an internal control. Each qPCR was carried out with 2 μL of cDNA or gDNA in 20 μL final volume following gene-specific amplification programs. The specificity of SYBR Green based qPCR assays was verified by the melting temperature (Tm) of the amplicon as calculated by the instrument software. Results were validated only when threshold cycle (Ct) values were under the limit value of 40 cycles and with an acceptable reproducibility between qPCR replicates (less than 5% of variation) (28).

### Histopathology

Kidney tissues were fixed in formalin, embedded in paraffin, sectioned and stained with hematoxylin & eosin (H&E). The histopathology was empirically quantified by scoring interstitial nephritis and fibrosis under a bright light microscope (29).

### Measurement of antibodies and creatinine in blood

Total IgG and IgG isotypes specific to *L. interrogans*, as well as creatinine present in serum was measured by ELISA using anti-Hamster IgG H+L, IgG1, IgG2 (Southern Biotech), and a creatinine assay kit (Sigma-Aldrich). For IgG determination whole cell sonicate of *L. interrogans* ser. Copenhageni FioCruz was used as the antigen.

### Statistics tests

Comparisons between male and female groups were analyzed by Mann-Whitney U Exact test (Fig. 1, Fig. 4, Fig. 5 and Fig. 6B). For histopathology scoring (Fig. 6A), one-tailed Fisher test was used. Differences in survival among experimental groups (Fig. 2) was determined using Logrank Exact test in R. Data analysis was done using GraphPad Prism 7 and QuickCals software. Statistical significance was set to *P* < 0.05.

**Figure 1.**
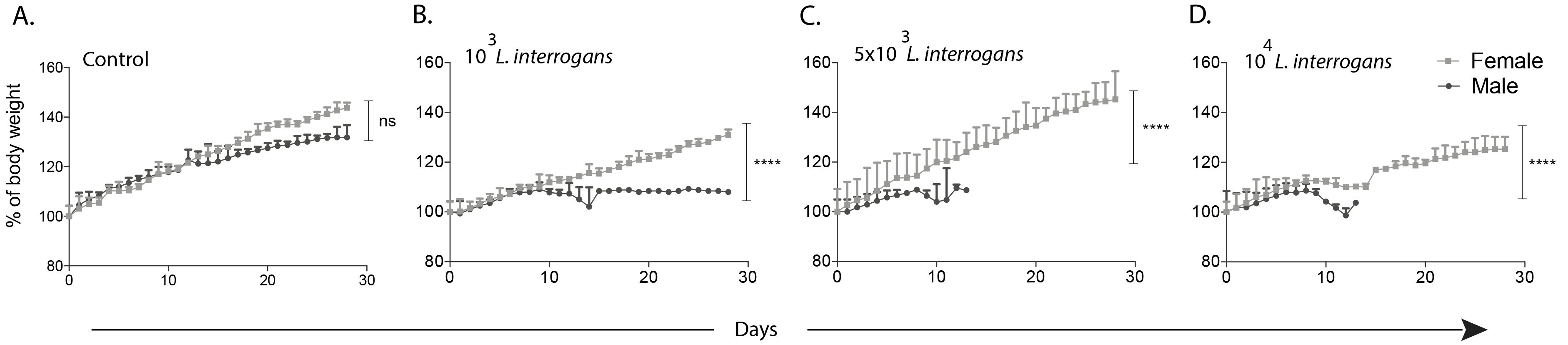
Body weight of male and female Golden Syrian hamsters infected with *L. interrogans*. Groups of hamsters were infected intraperitoneally with increasing doses of *L. interrogans* serovar Copenhageni strain Fiocruz L1-130. Body weight measurements (g) were recorded for 28 days post-infection and normalized to 100% on day 0 of infection (d0). Asterisks indicate significant differences between males and females by Mann-Whitney U Exact test, **** p < 0.0001; ns, not significant p=0.2873. Number of animals: 4 animals per group per sex (total n=32 hamsters). Data is representative of two experiments.

## RESULTS

### Female hamsters are more resistant to Leptospirosis than their male counterparts

Groups of hamsters were infected intraperitoneally with 10^3^, 5×10^3^ or 10^4^ *L. interrogans* serovar Copenhageni strain Fiocruz L1-130 and were monitored for objective clinical signs of infection (weighloss and survival) for 28 days. Overall, female hamsters gained body weight significantly (between 20 - 40%), whereas male survivors increased body weight by only 8% (p<0.0001); differences in weight were also observed in non-infected controls although the latter were not significant (Fig. 1). Regarding survival, female hamsters infected with 10^3^ or 5×10^3^, did not develop any symptom of disease and 100% of these survived, whereas only 25% and 0% of the groups of males infected with the same doses survived, respectively (Logrank Exact test p=0.1429 and p=0.0285); in the groups infected with 10^4^ bacteria, 37.5% of females survived whereas 0% of the males infected with the same dose survived, p×0.0085 (Fig. 2). Non-survivors reached endpoint criteria between days 9-15 post-infection. Two male hamsters succumbed to *Leptospira* infection before showing signs of disease or reaching weight loss endpoint criteria. All the non-infected controls (n=8) survived 28 days until termination. Overall, 15/16 (93.7%) male hamsters succumbed to infection whereas 2/16 (12.5%) female hamsters succumbed to the same infection doses. Overall differences between all the infected males and females (10^3^, 5×10^3^, and 10^4^) was analyzed with Logrank Exact test and it was significant, p<0.0001.

**Figure 2.**
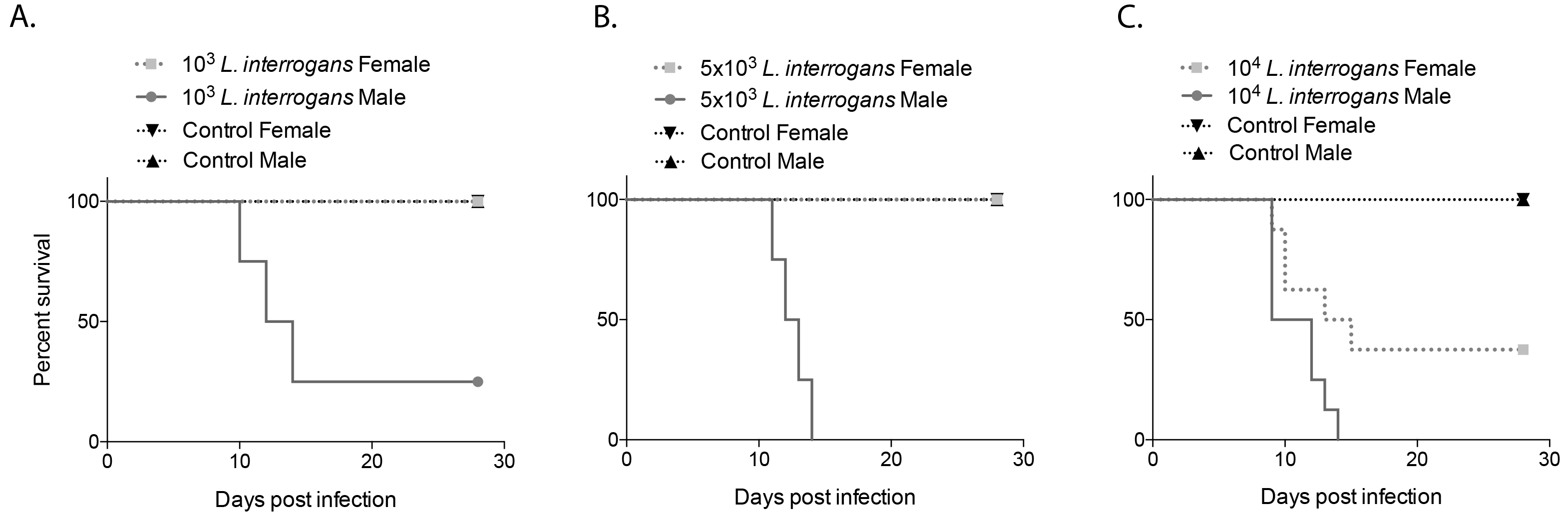
Percent survival of male and female hamsters after infection with increasing doses of *L. interrogans*. Groups of male and female hamsters were infected intraperitoneally with 10^3^ (A), 5×10^3^ (B), or 10^4^ (C) *L. interrogans* and with PBS (control) on day 0 and clinical scores were monitored for 28 days. Hamsters that reached endpoint criteria were euthanized. Number of hamsters: 10^3^, 5×10^3^ infections and controls, n=4 per group (A and B analysis, n=16 hamsters each); 10^4^ infection n=8 per group, controls n= 4 per group (C analysis, n=24 hamsters). Statistics by Logrank Exact test in R: A p=0.1429, B p=0.0285, C p=0.0085. A+B+C Logrank Exact Test in R p< 0.0001. Data is representative of three experiments.

### Leptospira burden was higher in kidney of male hamsters

The kidneys of male and female hamsters were collected at termination and the presence of leptospiral DNA was detected by PCR. Male hamsters had significantly higher (2-5 logs) genome equivalents of *L. interrogans* in the kidney than the females in the respective infected groups; infection with the lowest dose (10^3^) resulted in a 4Log increase of *L. interrogans* in the kidneys of male hamsters 28 days later, whereas the females had a reduction of ~1Log (Fig. 3). In males, increasing doses did not lead to increased amounts of *Leptospira* in the kidney (remained around 10^7^/mg tissue), whereas the opposite was true for females that showed an exponential increase of *Leptospira* in kidney as a result of increased infection doses (~10^2^ to 10^6^/mg tissue) (Fig. 3). The viability of *L. interrogans* in the kidney was analyzed by dark field microscopy of EMJH cultures of kidney tissue collected at termination. Cultures were checked for the presence of bacteria after 30 days of incubation, 91.6% of the infected males were culture positive and 66.6% of the females were positive for *L. interrogans* which shows a 25% difference in bacterial clearance from the kidney depending on sex.

**Figure 3.**
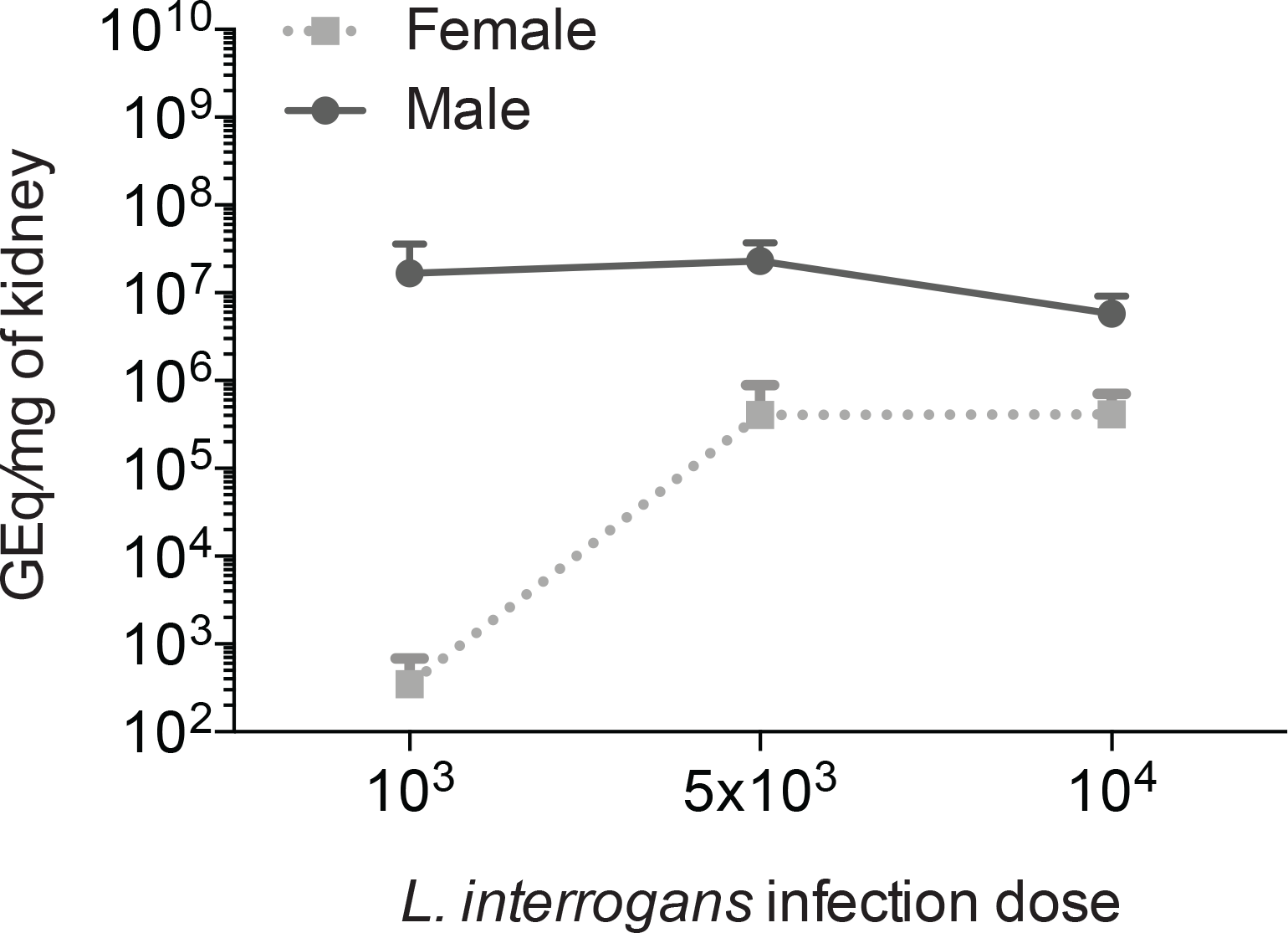
Quantification of *L. interrogans* DNA in kidney tissue of infected male and female hamsters. DNA was quantified by real-time PCR targeted to the *Leptospira* 16S rDNA purified from kidney from infected animals (n=23). Data is representative of two experiments. GEq, genome equivalents.

### Male hamsters had signs of kidney function impairment

Blood collected from hamsters at termination was used to determine the amount of total IgG, IgG1 and IgG2 antibodies and the concentration of creatinine in serum, by ELISA. No significant difference in IgG response was observed between males and females in the infected groups (OD_450_ Avg ± Stdev: females, 3.407 ± 0.485; males, 3.765 ± 0.124); IgG isotyping showed that levels of IgG2 were 2.5-fold higher than IgG1 without differences between sexes. Levels of creatinine (Fig. 4) were 2-fold greater in serum from male hamsters (5.71 ng/ul) than in females (2.67 ng/ul). Increased levels of creatinine in blood is a sign that glomerular filtration in the kidney is compromised.

**Figure 4.**
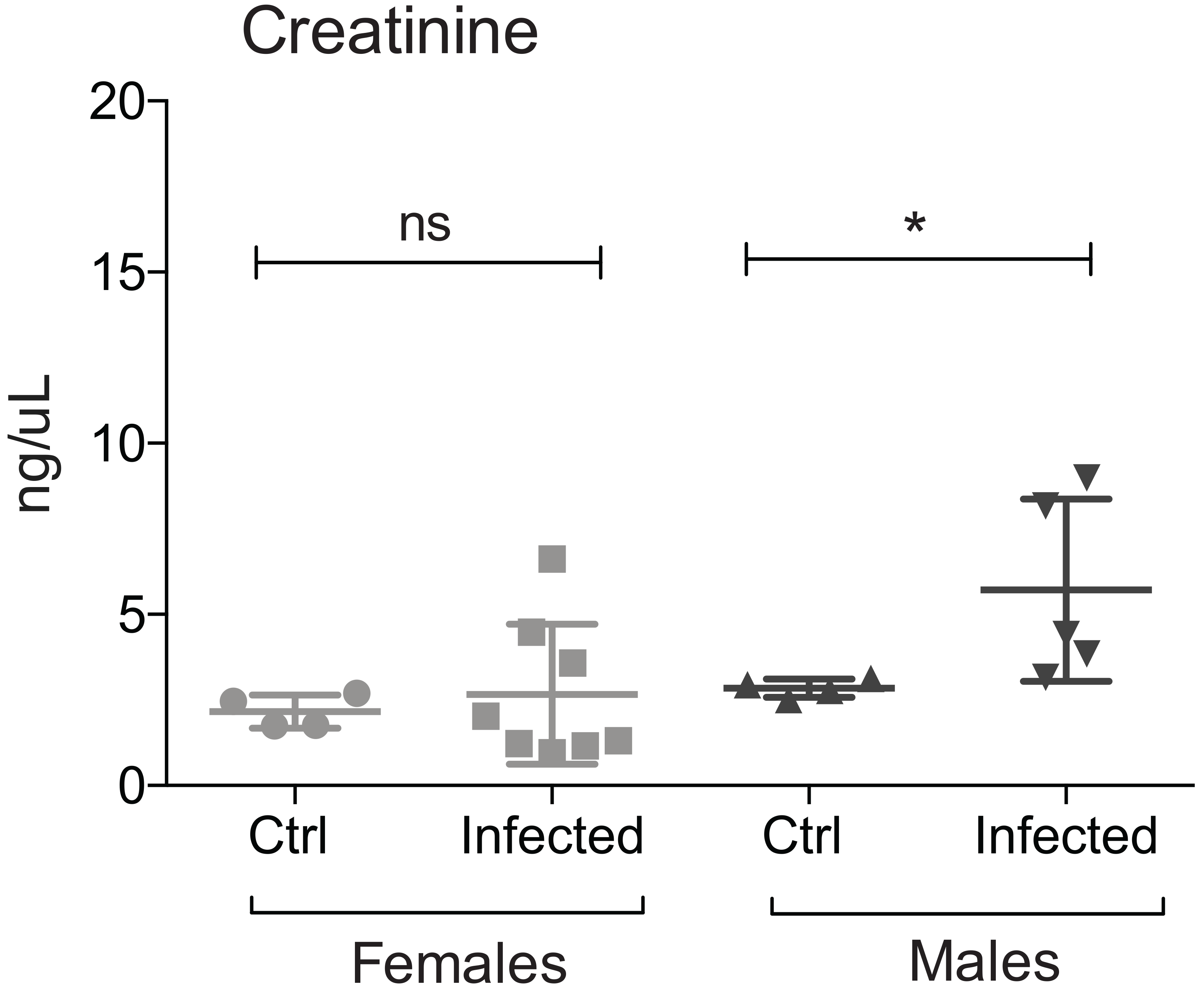
Kidney function of infected and non-infected hamsters. The concentration of creatinine was measured in blood collected at termination from male and female hamsters (n=21) infected with *L. interrogans* as well as from the respective controls. Statistical analysis by Mann Whitney U Exact test, ns, p=0.8081, * p=0.0159. Data is representative of two experiments.

### Chemokine and cytokine gene expression is higher in kidney of non-survivors than in kidney of survivor hamsters infected with *L. interrogans*

Gene expression of six pro-inflammatory markers CCL3/MIP-1α, CxCL10/IP-10, TNF-α, INF-γ, IL-4, and IL-6 and anti-inflammatory IL-10 were quantified in kidney of hamsters infected with 10^3^, 5×10^3^ or 10^4^ *L. interrogans* by RT-qPCR (Fig. 5). We compared the expression profiles of inflammatory markers between hamsters that met endpoint criteria before day 28 (non-survivors: 14 male, 5 female, n=19, 74% male), hamsters that reached endpoint at d28 (survivors: 1 male, 10 females, n=11, 91% female) and controls (4 male, 4 female, n=8). All three innate response pro-inflammatory markers CCL3/MIP-1α, CxCL10/IP-10 and TNF-α were significantly up-regulated in non-survivor kidneys compared to survivor tissue (Fig. 5A), whereas adaptive pro-inflammatory cytokines INF-γ, IL-4 and IL-6 were not significantly different between non-survivors and survivors (Fig. 5B). The expression profile of anti-inflammatory IL-10 was significantly increased in the non-survivor group (Fig. 5C).

**Figure 5.**
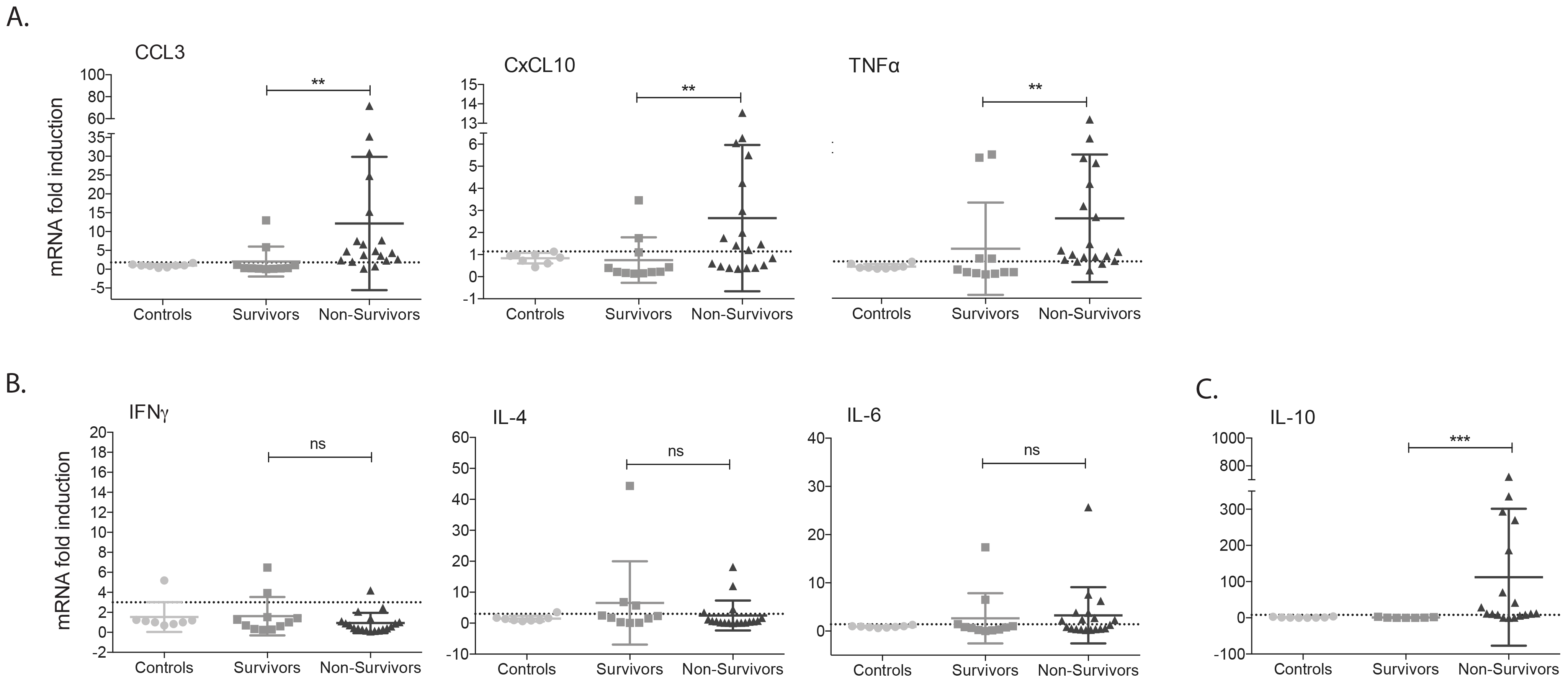
Quantification of inflammatory chemokines and cytokines in kidney of survivor and non-survivor hamsters. The gene expression in the kidney of hamsters infected with *L. interrogans* serovar Copenhageni was quantified using real-time PCR (2-ΔΔCt method) and β-actin mRNA as an internal standard. A. Innate immune response CCL3/MIP-1α, CxCL10/IP10 and TNFα; B. Adaptive immune response IFN-γ, IL-4, and IL-6; and anti-inflammatory IL-10. Statistical analysis by Mann-Whitney Exact test (CCL3 p=0.0017, CxCL10 p=0.0045, TNFα p=0.0094, IFNγ p=0.2497, IL-4 p=0.2449, IL-6 p=0.3495 and IL-10 p=0.0005). Number of hamsters, n=38. Data is representative of three experiments.

### Kidney of infected non-survivor hamsters had higher histopathological scores and more fibrosis biomarker ColA1 than kidney from survivors

Histopathological analysis of kidneys from infected hamsters stained with H&E showed that non-survivors had a significant increase in interstitial nephritis with infiltrates of mononuclear cells (~58%) and increased interstitial collagen deposition (~33%) when compared with survivor hamsters (~15% and 7%, respectively) (Fig 6A). At the molecular level, fibroblast activation marker (collagen1 A1, ColA1) mRNA expression was 10× higher in kidneys of infected non-survivors (2.7 ± 2.5) than in survivors (0.27 ± 0.14). However, inducible nitric oxide (iNOS) which has been associated both with potentially resolving inflammation as well as fibrosis in mouse kidney, was not different between survivor and non-survivor groups (Fig. 6B).

**Figure 6.**
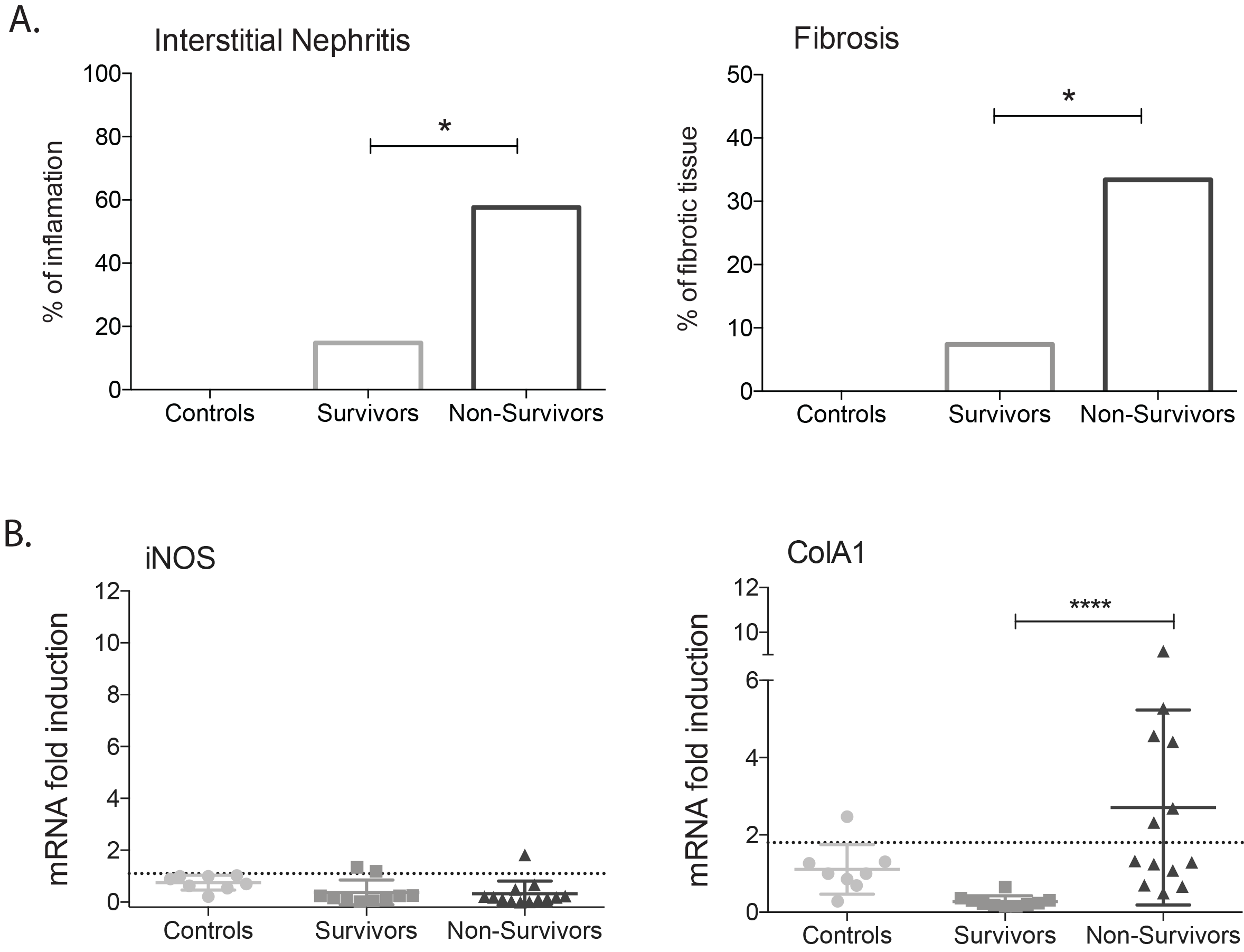
Histopathological scores and quantification of inflammation and fibrosis markers in kidney of survivor and non-survivor hamsters. A, Histopathology was empirically quantified by scoring interstitial nephritis and fibrosis in kidney sections of individual hamsters (n=20) infected and not infected with *L. interrogans*; statistics by Fisher one-tailed test, * p < 0.05; B, mRNA gene expression of iNOS and ColA1 in kidney (n=31). Statistics by Mann-Whitney U Exact test, ColA1 p<0.0001. Data is representative of two experiments.

## DISCUSSION

There is evidence from clinical and seroepidemiologic data that sex impacts severity of leptospirosis with a bias toward higher hospitalization rates for men (8), (9), (10), (11), (12), (13). A comprehensive study of sex differences in clinical leptospirosis in Germany found that male patients were more likely to be hospitalized than female patients and suggested that reports on male predominance in leptospirosis may thus reflect sex-related variability in the incidence of severe disease rather different infection rates (30). However, several reports of leptospirosis outbreaks where males and females have similar levels of exposure, have found no significant effects of sex differences on development of illness (31), (32), (33).

Very few researchers have devoted resources to evaluate the question of susceptibility to severe disease related to sex in animal models. We evaluated how sex affects pathology, disease progression and mortality after *Leptospira* infection using an acute model of leptospirosis and we found that male hamsters are considerably more susceptible to lethal infection with pathogenic Leptospira than female animals: 93.7% of males did not survive infection as opposed to 87.5% of females who did survive the same infectious doses. Analysis of weight of the animals infected with increasing doses of *Leptospira* showed that males are more susceptible to lose weight as a sign of disease progression and to meet endpoint criteria sooner than females exposed to the same conditions, independently of the infecting dose. These data are consistent with another comparative study that assessed severity of pulmonary leptospirosis in female and male hamsters. In that study, male hamsters developed pulmonary hemorrhage after infection with *L. interrogans* serovar Hebdomadis 120 h post-infection, whereas its female counterparts did not (34). One study of human leptospirosis performed in the Netherlands, showed that 90% of the reported cases happened in male patients and that the severity of the disease among males does not appear to be related to infections caused by more virulent serovars (12). However, in a study performed in Germany, Jansen et al found that male patients (n=263) had more severe clinical outcomes of leptospirosis and higher case fatality rates relative to their female counterparts (n=75), (5% versus 1%, respectively) without significant differences in the type of exposure, or time from onset of symptoms to treatment (4.5 days for both) (30). Even though it may be difficult to resolve whether the disease is aggravated by genetic, hormonal, and immunological differences or to exposure-related or to behavior-related issues associated with male behavior (35), our results corroborate Tomizawa (hamster) and Jansen’s (human) findings and suggest that female hamsters, exposed to the same
infectious conditions as males, are more resistant to development of symptoms and develop milder disease. In this case, regarding exposure to the same infectious doses under the same conditions, sex differences appear to play an important role in the severity of leptospirosis.

Another observation from our study is that male hamsters were equally susceptible to low and high doses of infection, whereas female susceptibility increased with higher infection doses. When we quantified the Leptospira DNA in the kidney, male hamsters infected with 10^3^ to 10^4^ *Leptospira* had a consistent high number of bacteria in the kidney (~ 10^6^ to 10^7^ per mg of tissue), whereas Leptospira DNA quantified in kidney from female hamsters grew exponentially with increased infectious doses (~10^2^ to 10^5^ bacteria per mg of tissue). Kidney culture also showed a higher recovery rate of viable *L. interrogans* in the kidney of males (91.6%) than females (66.6%). We speculate that females may be able to mount a more effective immune response to lower doses of *Leptospira* but when the infectious dose reaches a certain threshold, colonization of the kidney takes place and the disease progresses with similar lethal results as in males.

Quantification of the inflammatory markers in kidney of non-survivors (mostly male) showed higher profiles of innate pro-inflammatory activity (CCL3, CxCL10 and TNF-α) in contrast to the survivors (mostly female) that had low expression profiles of these markers. Higher expression levels of CCL3, CxCL10 and TNF-α correlated with more interstitial nephritis and fibrosis in kidneys, and with worse survival outcomes. In addition, anti-inflammatory IL-10 was significantly increased in kidneys of non-survivors. Adaptive immune response cytokine expression, IFNg, IL-4 and IL-6, were not different between survivor and non-survivor groups which suggests that TH1, TH2 and Treg responses may have not been engaged. Our results corroborate previous observations in humans, where patients with severe leptospirosis experience a cytokine storm characterized by high levels of TNF-α and IL-10 (36), (2). Similarly, high levels of TNF-α, IL-1beta, IL-6, IL-8 and IL-10, and were reported in serum from septic patients (37). Also, another report showed higher levels of CxCL10 (IP-10) which is associated with cell-mediated immunity in leptospirosis patients (38). These data suggest that for *Leptospira* eradication, less pro-inflammatory response correlate with better pathology and survival outcomes. Production of pro-inflammatory chemokines reflects an immune response that may promote further renal damage. Fibroblast activation marker (collagen1 A1, ColA1) and iNOS are known to play important roles in *Leptospira-induced* interstitial nephritis (39), (40) and kidney fibrosis (41) in mice. The quantification of mRNA transcripts in kidney of non-survivor hamsters showed up-regulation of ColA1, but not iNOS, which was absent. These results suggest that, in hamsters, ColA1 is a marker of severe kidney pathology.

Our results indicate that male hamsters infected with *L. interrogans* ser. Copenhageni succumb to severe leptospirosis after exposure to infection doses 1Log lower than females. Once the critical dose is reached females also succumb to infection. It remains to be seen if this difference in susceptibility can be explained by distinctions in immune responses to pathogenic *Leptospira* between males and females that may account for more effective control of *L. interrogans* dissemination in females early in the infection. In endemic areas, humans are more often exposed to the lower infection doses (~10^3^) (42) which were lethal to males, than the higher doses (>10^4^) which were lethal to males and females. Thus, it is possible that the frequency of lower infectious doses in nature and increased biological susceptibility both contribute for aggravated outcomes of leptospirosis in males. It is notable that most studies using acute animal models of infection have been done using males. Consequently, there’s a male-bias in the knowledge acquired for Leptospirosis to date. Lethal leptospirosis needs to be further explored in side-by-side comparative studies between males and females in hamsters and in other animal models.

## Acknowledgments

We thank the Research Histology Core of UTHSC for providing the histopathology scores and the staff at the UTHSC BERD (Biostats, Epidemiology and Research Design) clinic for support with statistical analysis. This work was supported by Public Health Service grants R44AI096551 (M.G.S.) and R43AI136551 (M.G.S.) from the National Institutes of Health.

